# Evaluation of the accuracy of imputed sequence variants and their utility for causal variant detection in cattle

**DOI:** 10.1101/085399

**Authors:** Hubert Pausch, Iona M MacLeod, Ruedi Fries, Reiner Emmerling, Phil J Bowman, Hans D Daetwyler, Michael E Goddard

## Abstract

**Background:** The availability of dense genotypes and whole-genome sequence variants from various sources offers the opportunity to compile large data sets consisting of tens of thousands of individuals with genotypes at millions of polymorphic sites that may enhance the power of genomic analyses. The imputation of missing genotypes ensures that all individuals have genotypes for a shared set of variants.

**Results:** We evaluated the accuracy of imputation from dense genotypes to whole-genome sequence variants in 249 Fleckvieh and 450 Holstein cattle using *Minimac* and *FImpute*. The sequence variants of a subset of the animals were reduced to the variants that were included in the Illumina BovineHD genotyping array and subsequently inferred *in silico* using either within-or multi-breed reference populations. The accuracy of imputation varied considerably across chromosomes and dropped at regions where the bovine genome contains segmental duplications. Depending on the imputation strategy, the correlation between imputed and true genotypes ranged from 0.898 to 0.952. The accuracy of imputation was higher with *Minimac* than *FImpute* particularly for variants with low MAF. Considering a multi-breed reference population increased the accuracy of imputation, particularly when *FImpute* was used to infer genotypes. When the sequence variants were imputed using *Minimac*, the true genotypes were more correlated to predicted allele dosages than best-guess genotypes. The computing costs to impute 23,256,743 sequence variants in 6958 animals were ten-fold higher with *Minimac* than *FImpute*. Association studies with imputed sequence variants revealed seven quantitative trait loci (QTL) for milk fat percentage. Two causal mutations in the *DGAT1* and *GHR* genes were the most significantly associated variants at two QTL on chromosomes 14 and 20 when *Minimac* was used to infer genotypes.

**Conclusions:** The population-based imputation of millions of sequence variants in large cohorts is computationally feasible and provides accurate genotypes. However, the accuracy of imputation is low at regions where the genome contains large segmental duplications or the coverage with array-derived SNPs is poor. Using a reference population that includes individuals from many breeds increases the accuracy of imputation particularly at low-frequency variants. Considering allele dosages rather than best-guess genotypes as explanatory variables is advantageous to detect causal mutations in association studies with imputed sequence variants.

## Background

Several genotyping arrays that comprise a varying number of single nucleotide polymorphisms (SNPs) are routinely used for genome-wide genotyping in cattle. Cows are usually genotyped using low-density genotyping arrays whereas bulls are mostly genotyped at higher density [1]. Moreover, the sequencing of important ancestors of many cattle breeds yielded genotypes at millions of polymorphic sites [2]. Combining the genotype data from various sources into a single large data set may enhance the power of genome-wide analyses. The imputation of missing genotypes is necessary to ensure that all individuals have genotypes for a shared set of variants. Genotype imputation may also infer dense genotypes *in silico* for animals that were genotyped at lower density using animals that were genotyped at a higher density as a reference [3].

Algorithms that infer missing genotypes apply family-(*e.g*., [4]) or population-based (*e.g*., [5–7]) imputation approaches or a combination thereof (*e.g*., [8–10]). Family-based imputation approaches rely on Mendelian transmission rules in pedigrees to infer missing genotypes. Population-based imputation approaches exploit linkage disequilibrium (LD) between adjacent markers to predict missing genotypes using a probabilistic framework without (explicitly) considering pedigree information [11]. While population-based imputation approaches are accurate, their computing costs are too high to infer genotypes for a large number of animals and markers for routine applications [12, 13]. Methods that apply a combination of family-and population-based imputation approaches exploit shared haplotypes among relatives thereby enabling rapid imputation of genotypes for tens of thousands of individuals and millions of markers *in silico* [8–10, 14].

The accuracy of imputation from low to higher density depends on the relationships between target and reference animals, genotype density in both panels, allele frequencies of the imputed variants, LD between adjacent markers and algorithms applied to infer missing genotypes (*e.g*., [12, 15, 16]). These parameters also affect the accuracy of imputation from dense genotypes to sequence variants [2, 17, 18]. However, the accurate imputation of low-frequency variants is critical with sequence data because rare alleles are more frequent in sequence than array-derived variants and the LD between low-frequency and array-derived variants may be low [19, 20].

A number of studies with simulated and real sequence data in cattle indicated that using imputed sequence variants may improve genomic predictions and facilitate the detection of causal trait variants because the polymorphisms that underlie phenotypic variation are included in the data [2, 19, 21–23]. However, the benefits of dense marker maps for genome-wide analyses may be compromised when the accuracy of imputation is low [24–26].

In this paper, we evaluate the accuracy of imputation for sequence variants in two cattle breeds using different imputation algorithms and reference populations. We perform association studies between imputed sequence variants and milk fat percentage in 6958 Fleckvieh bulls and show that the imputation strategy is critical to pinpoint causal mutations.

## Methods

### Animal ethics statement

No ethical approval was required for this study.

### Sequenced animals

We used whole-genome sequence data of 1577 taurine animals that were included in the fifth run of the of the 1000 bull genomes project [2]. The reads were aligned to the UMD3.1 bovine reference genome using the *BWA-MEM* algorithm [27, 28]. SNPs, short insertions and deletions were genotyped for all sequenced animals simultaneously using a multi-sample variant calling pipeline that was implemented with the *mpileup* module of *SAMtools* [29] and that is described in Daetwyler et al. [2]. The variant calling yielded genotypes at 39,721,987 biallelic sites for 1577 animals. We considered genotypes at 22,737,136 autosomal sequence variants that had a minor allele frequency (MAF) higher than 0.5% to build the genomic relationship matrix among the sequenced animals using the *plink* (version 1.9) software tool [30]. Principal components of the genomic relationship matrix were calculated using the *GCTA* (version 1.25.3) software tool [31]. Our analyses focused on the Fleckvieh (FV) and Holstein (HOL) breeds because a large number of animals from both breeds were sequenced for the 1000 bull genomes project. Following the inspection of the top principal components, animals whose breed was uncertain were removed leaving 249 FV and 450 HOL cattle (see **Additional file 1 Figure S1**).

### Design of cross-validation scenarios

The imputation from dense genotypes to full sequence variants was evaluated using 15-fold cross-validation in FV and HOL cattle. We considered only sequence variants that were polymorphic in 249 FV or 450 HOL animals. For each fold, the sequenced animals were divided into reference and validation animals. Forty-nine FV and 100 HOL animals, that were a random subset of the sequenced animals, were considered as validation animals (*i.e*., the same individual may have been included in several validation sets). All remaining sequenced animals from the 1000 bull genomes project were considered as a multi-breed reference population. The within-breed reference populations consisted of 200 FV or 350 HOL animals. The genotypes of the validation animals were reduced to the variants that were included in the Illumina BovineHD genotyping array (HD) to mimic dense genotypes. The masked genotypes of the validation animals were subsequently inferred *in silico* using full sequence information from the reference animals. The selection of validation animals and subsequent imputation of genotypes was repeated 15 times for chromosomes 1, 5, 10, 15, 20 and 25.

### Evaluation of the accuracy of imputation

The overall accuracy of imputation was the mean correlation between *in silico* imputed and true (sequenced) genotypes (r_IMP,SEQ_) across 15 folds and six chromosomes analysed. Specifically, for each fold and chromosome, we calculated the correlation between matrices of imputed and true genotypes. These values were averaged to obtain the overall r_IMP,SEQ_. We additionally grouped the imputed sequence variants into 50 classes with regard to their MAF in the reference population. The r_IMP,SEQ_-value for each MAF class was the mean value across six chromosomes and 15 folds. To detect intra-chromosomal variations in the accuracy of imputation, we calculated r_IMP,SEQ_-values for successive 1 Mb segments along each chromosome analysed. Differences between imputation scenarios were tested for significance using two-tailed t-tests.

### Imputation methods

The performance of the *FImpute* (version 2.2) [9] and *Minimac3* (version 2.0.1, henceforth referred to as *Minimac*) [7] software tools was evaluated using default parameter settings. The algorithm implemented in *FImpute* uses family and population-based information to infer haplotypes and missing genotypes. A pedigree consisting of 47,012 FV animals tracing back ancestry information to animals born in 1925 was considered when genotypes were inferred using *FImpute*. The pedigree included ancestry information for 138 (out of 250) sequenced FV animals. Pedigree records for sequenced animals from other breeds were also included if available. However, these pedigrees included only first-degree relatives. Genotypes for the imputed sequence variants were coded as 0, 1 and 2 for homozygous, heterozygous and alternative homozygous animals, respectively.

The algorithm implemented in the *Minimac* software tool uses previously phased genotypes, *i.e*., it takes haplotypes as input for reference and validation animals. Haplotype phases were estimated separately for the reference and validation animals using the phasing algorithm implemented in the *Eagle* (version 2.3) software tool [32]. Both *Eagle* and *Minimac* do not consider pedigree information to infer haplotypes and missing genotypes. *Minimac* provides best-guess genotypes (coded as 0, 1 and 2 for homozygous, heterozygous and alternative homozygous animals, respectively) and allele dosages (continuously distributed values ranging from 0 to 2) for the imputed sequence variants. The r_IMP,SEQ_-values were calculated for best-guess genotypes (*Minimac*_BG_) and allele dosages (*Minimac*_DOS_).

### Imputation of sequence variants in 6958 Fleckvieh animals

We imputed genotypes for 23,256,743 autosomal sequence variants that were polymorphic in 249 sequenced FV animals for 6958 FV bulls that had (partially imputed) array-derived genotypes at 603,662 autosomal variants (see [33]). Genotypes were imputed with either *FImpute* or *Minimac* using either within-or multi-breed reference populations that consisted either of 249 FV animals or 1577 animals from various bovine breeds (see above). Haplotype phases for reference and target animals were estimated using the *Eagle* software tool (see above). The LD among imputed sequence variants was calculated as the squared correlation between predicted allele dosages.

### Accuracy of imputation at two known causal mutations

712 and 902 FV animals that had imputed sequence variants also had direct genotypes for two mutations in the *DGAT1* (rs109234250 and rs109326954, p.A232K) and *GHR* (rs385640152, p.F279Y) genes that were obtained for previous studies using TaqMan^®^ genotyping assays (Life Technologies) [34–37]. The accuracy of imputation at both sites was the proportion of correctly imputed genotypes.

### Association analyses in Fleckvieh cattle

Association tests between 23,256,743 imputed sequence variants and milk fat percentage were carried out in 6958 FV bulls using a variance components-based approach that was implemented in the *EMMAX* software tool and that accounts for population stratification by fitting a genomic relationship matrix as detailed in [38] and [20]. The genomic relationship matrix was built using array-derived genotypes of 603,662 (partially imputed) autosomal SNPs [39]. Daughter yield deviations (DYDs) for milk fat percentage were the response variables. The genotypes were coded as 0, 1 and 2 for homozygous, heterozygous and alternative homozygous animals, respectively, when sequence variants were imputed using *FImpute*. When the sequence variants were imputed using *Minimac*, we considered both best-guess genotypes and allele dosages as explanatory variables for the association tests. Sequence variants with P values less than 2.1×10^−9^ (5%-Bonferroni-corrected type I error threshold for 23,256,743 tests) were considered as significantly associated.

### Computing environment

All computations were performed on the Biosciences Advanced Scientific Computing (BASC) cluster that is located at AgriBio, Centre for AgriBiosciences, VIC 3083, Bundoora. The memory usage and process time required to infer genotypes for 23,256,743 sequence variants in 6958 animals was quantified on 12-core Intel^®^ Xeon^®^ processors rated at 2.93 Ghz with 96 gigabytes of random-access memory (RAM).

## Results

We evaluated the accuracy of imputation from dense genotypes to sequence variants in 249 FV and 450 HOL animals using sequence data on bovine chromosomes (BTA for Bos taurus) 1, 5, 10, 15, 20 and 25. The number of polymorphic sites ranged from 413,371 (BTA25) to 1,444,299 (BTA1) and from 383,072 (BTA25) to 1,382,987 (BTA1) in FV and HOL, respectively, indicating that genetic diversity is higher in FV than HOL cattle (Table 1). Variants with low MAF were more frequent among the sequence than HD variants; between 58.12 and 60.55% of the sequence variants and between 14.27 and 18.55% of the HD variants had a MAF lower than 10%.

**Table 1.** Number of polymorphic sites in Fleckvieh and Holstein cattle

| Chr | Chr.<br>Length<br>(Mb) | Fleckvieh |  | Holstein |  |
| --- | --- | --- | --- | --- | --- |
|  |  | SEQ (MAF<0.1) | 700K (MAF<0.1) | SEQ (MAF<0.1) | 700K (MAF<0.1) |
| 1 | 158.32 | 1,444,299 (60.55) | 38,009 (18.40) | 1,382,987 (59.11) | 37,397 (17.01) |
| 5 | 121.18 | 1,098,976 (59.64) | 28,173 (18.55) | 1,073,964 (59.12) | 27,667 (15.90) |
| 10 | 104.30 | 959,206 (59.13) | 25,787 (17.09) | 917,799 (59.48) | 25,497 (16.12) |
| 15 | 85.27 | 866,151 (58.54) | 20,617 (17.24) | 827,526 (58.12) | 20,360 (14.38) |
| 20 | 71.98 | 679,738 (59.94) | 18,466 (18.18) | 649,768 (59.18) | 18,264 (18.13) |
| 25 | 42.85 | 413,371 (59.08) | 11,370 (14.69) | 383,072 (58.14) | 11,179 (14.27) |

Number of sequence (SEQ) and array-derived (Illumina bovineHD, 700K) variants that were polymorphic in 249 Fleckvieh and 450 Holstein animals. The proportion of variants with a minor allele frequency (MAF) lower than 10% is given in parentheses.

### Evaluation of the accuracy of imputation

The correlation between imputed and sequenced genotypes (r_IMP,SEQ_) was higher in HOL than FV cattle (Table 2) likely reflecting lower genetic diversity and larger reference population size in HOL. When within-breed reference populations were considered, the ri_MP_,_SE_Q-values were 0.898, 0.908 and 0.934 for *FImpute, *Minimac*_BG_* and *Minimac*_DOS_ in FV and 0.912, 0.929 and 0.951 in HOL, respectively. Adding animals from various breeds to the reference panel increased ri_M_p,sEQ in FV, particularly when *FImpute* was used to impute missing genotypes (P=7.5×10^−11^, Table 2). The highest accuracy of imputation in FV (r_IMP,SEQ_=0.939) was achieved using *Minimac*_DOS_ with a multi-breed reference population. In HOL, a multi-breed reference population increased r_IMP,SEQ_ when *FImpute* was used to infer missing genotypes (P=4.0×10^−4^). However, the use of a multi-breed reference population had little effect on the r_IMP,SEQ_-values in HOL when *Minimac* was used (P>0.12). Using *Minimac*_DOS_ and a within-breed reference population provided the highest accuracy of imputation (r_IMP,SEQ_=0.951) in HOL. Regardless of the reference population used, the r_IMP,SEQ_-values were higher for *Minimac*_DOS_ than *Minimac*_BG_ in both breeds (P_F_v<2.9×10^−6^, Phol<3.9×10-^6^).

**Table 2.** Cross-validation imputation accuracy in Fleckvieh and Holstein cattle

| Breed | Within-breed reference populations |  |  | Multi-breed reference populations |  |  |
| --- | --- | --- | --- | --- | --- | --- |
|  | *Fimpute* | *Minimac*<sub>BG</sub> | *Minimac*<sub>DOS</sub> | *Fimpute* | *Minimac*<sub>BG</sub> | *Minimac*<sub>DOS</sub> |
| Fleckvieh | 0.898 | 0.908 | 0.934 | 0.921 | 0.916 | 0.939 |
|  | (0.674) | (0.735) | (0.771) | (0.774) | (0.812) | (0.838) |
| Holstein | 0.912 | 0.929 | 0.951 | 0.926 | 0.928 | 0.948 |
|  | (0.681) | (0.772) | (0.798) | (0.732) | (0.797) | (0.820) |

The values represent the correlation between true and imputed genotypes. The correlation between true and imputed genotypes for sequence variants with a minor allele frequency lower than 10% is given in parentheses. BG: best-guess, DOS: allele dosage.

A decline in the accuracy of imputation was evident for low-frequency variants across all scenarios tested. While r_IMP,SEQ_ was high for variants with a MAF higher than 10%, it was considerably less for low MAF variants (Figure 1a-b). In FV, the accuracy of imputation was higher for low MAF variants using multi-than within-breed reference populations (P<6.6×10^−6^, Table 2). The benefit of multi-breed reference panels was less pronounced in HOL and it was only significant (P=0.013) when *FImpute* was used. The r_IMP,SEQ_-values for low-frequency variants (MAF<0.1) were higher with *Minimac* than *FImpute* (P_FV_<0.005, P_HOL_<0.002, Table 2, Figure 1a-b).

**Figure 1.**
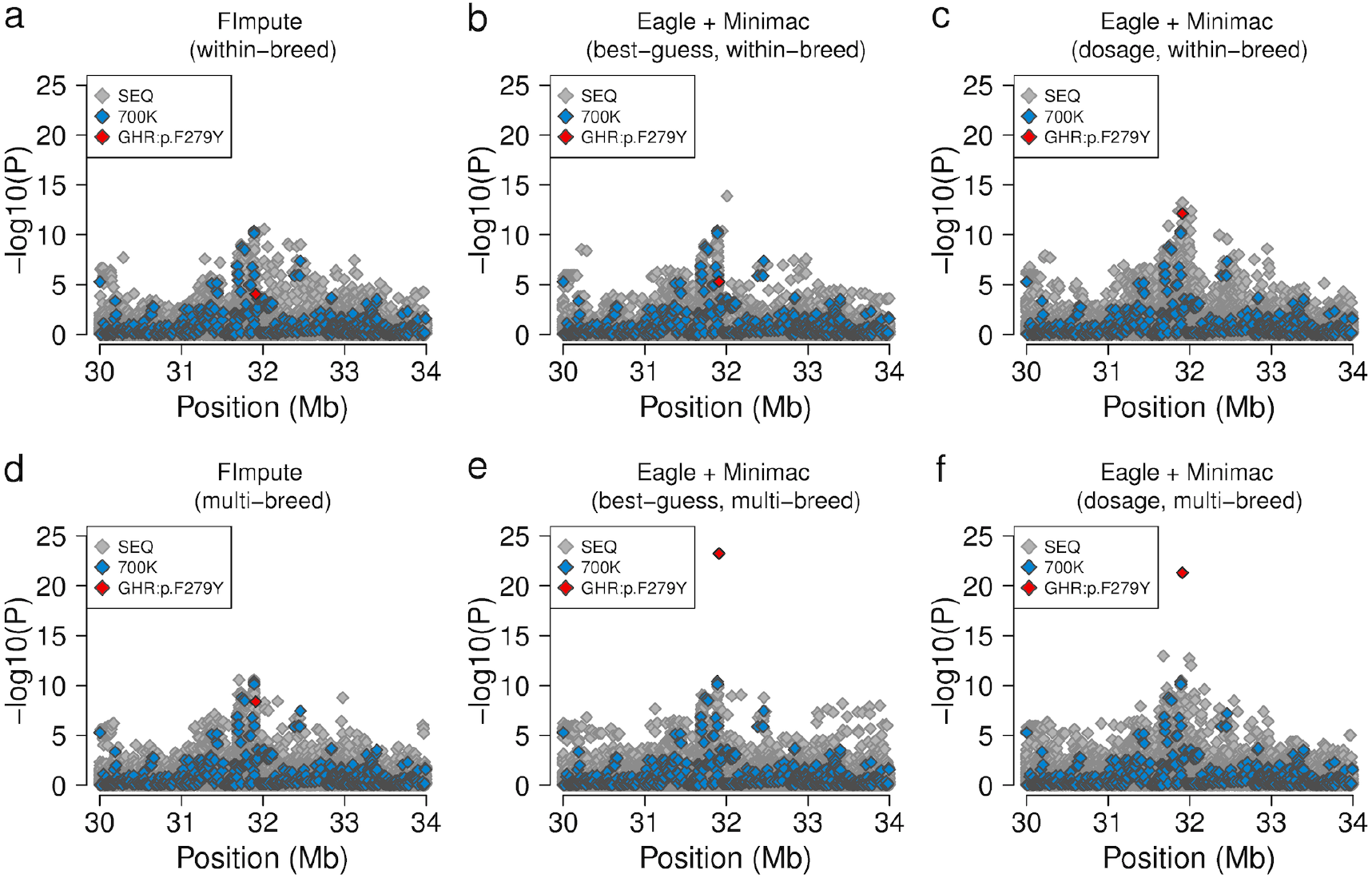
The accuracy of imputation in Fleckvieh and Holstein cattle. (a-b) The correlation between true and imputed genotypes as a function of the frequency of the minor allele in Fleckvieh (a) and Holstein (b) animals. (c-d) The boxplots represent the correlation between true and imputed genotypes for sequence variants on six chromosomes in Fleckvieh (c) and Holstein (d) animals.

The number of sequence variants that were not polymorphic after the imputation in the validation population varied considerably across the imputation scenarios tested (see **Additional File 2 Figure S2**). Using *Minimac*_HC_, between 29.2 and 37.4% of the imputed sequence variants were invariant in the validation populations depending on the breed and reference population considered. The proportion of invariant sites was lower when *FImpute* (between 19.8 and 24.2% of the sequence variants) or *Minimac*_DOS_ (between 3.6 and 6.7% of the sequence variants) were used for the imputation. Variants that had a low MAF in the reference population were more often invariant in the validation population than common variants. When the r_I_M_P_,sEQ-values were calculated using only sequence variants that were polymorphic in the validation population, *Minimac_HC_* provided the most accurate genotypes. However, the accuracy of imputation was also high using *Minimac*_DOS_ (see **Additional File 2 Figure S2c,d**) and it provided imputed genotypes for considerably more sequence variants than *Minimac_HC_* and *FImpute*.

The accuracy of imputation varied considerably across chromosomes. The r_IMP,SEQ_-values were high for sequence variants located on BTA1, BTA20 and BTA25. However, the accuracy of imputation was lower for sequence variants located on BTA5, BTA10 and BTA15 (Figure 1c-d). This pattern was observed regardless of the reference population and imputation software used. To investigate the reason for the variable accuracy of imputation across chromosomes, we calculated r_IMP,SEQ_-values, HD SNP coverage and sequence variant density for successive 1 Mb windows along the six chromosomes analysed. Multiple (partially extended) segments with high imputation errors were located on chromosomes 5, 10 and 15 (see **Additional file 3 Figure S3**) at positions where the bovine genome contains large segmental duplications [40],[41]. These segments were often sparsely covered with HD SNPs but contained a large number of polymorphic sequence variants. Such segments with low imputation quality were not detected along chromosomes 1, 20 and 25, that had an overall higher accuracy of imputation.

### Pinpointing causal mutations among imputed sequence variants

We imputed genotypes for 756,135 and 679,738 sequence variants that were located on BTA14 and BTA20, respectively, in 6958 FV animals using either *FImpute* or *Minimac* and considering either within-or multi-breed reference populations. To evaluate the ability to detect causal mutations, the imputed genotypes were tested for association with milk fat percentage. Milk fat percentage was the target trait because two mutations in the *DGAT1* (rs109326954, pA232K) and *GHR* (rs385640152, p.F279Y) genes have large effects on the milk composition in many cattle breeds including FV. In 249 sequenced FV animals, the minor allele frequencies of rs109326954 and rs385640152 were 6.2 and 7.2%. Both variants were more frequent in the multi-breed reference population (17.3 and 12.1%, Table 3). Depending on the reference population and imputation algorithm used, the proportion of correctly imputed genotypes was between 99.3 and 99.9% and between 86.1 and 99.4% for rs109326954 and rs385640152, respectively (Table 3). The imputation error rates were lower with *Minimac* than *FImpute*. Using *Minimac* and a multi-breed reference population provided the most accurate genotypes at both variants (99.9 and 99.4%).

**Table 3.**
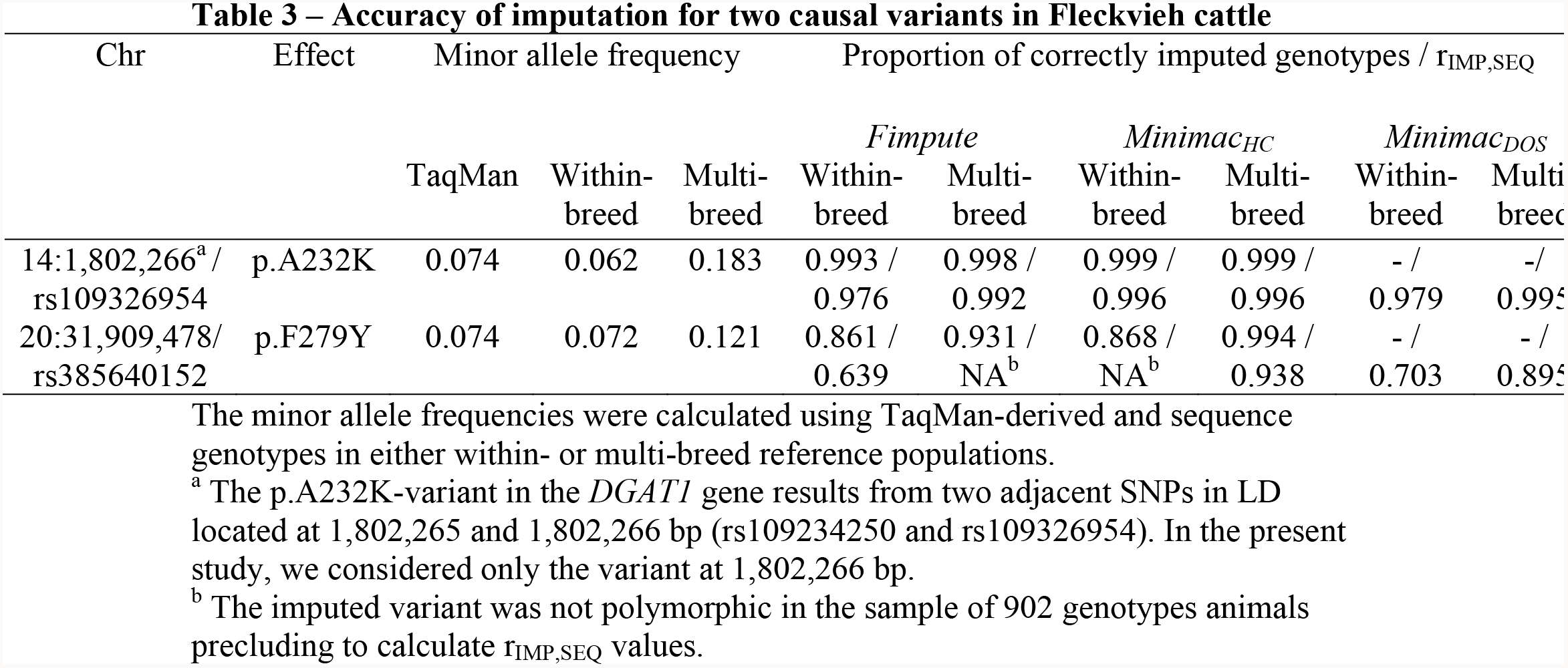
Accuracy of imputation for two causal variants in Fleckvieh cattle.

Association tests between the imputed genotypes and milk fat percentage revealed that the *DGAT1*:p.A232K-variant was among the most significantly associated variants across all scenarios tested reflecting high accuracy of imputation regardless of the reference population and imputation algorithm used (Figure 2, Table 3). However, several adjacent variants (±90 kb) were in complete or near complete LD (r^2^>0.99) with rs109326954 and had P values that were slightly higher, identical, or slightly lower. When the genotypes were imputed using *Minimac* and considering a within-breed reference population, five variants in complete LD including rs109326954 were the most significantly associated (Figure 2e,f).

**Figure 2.**
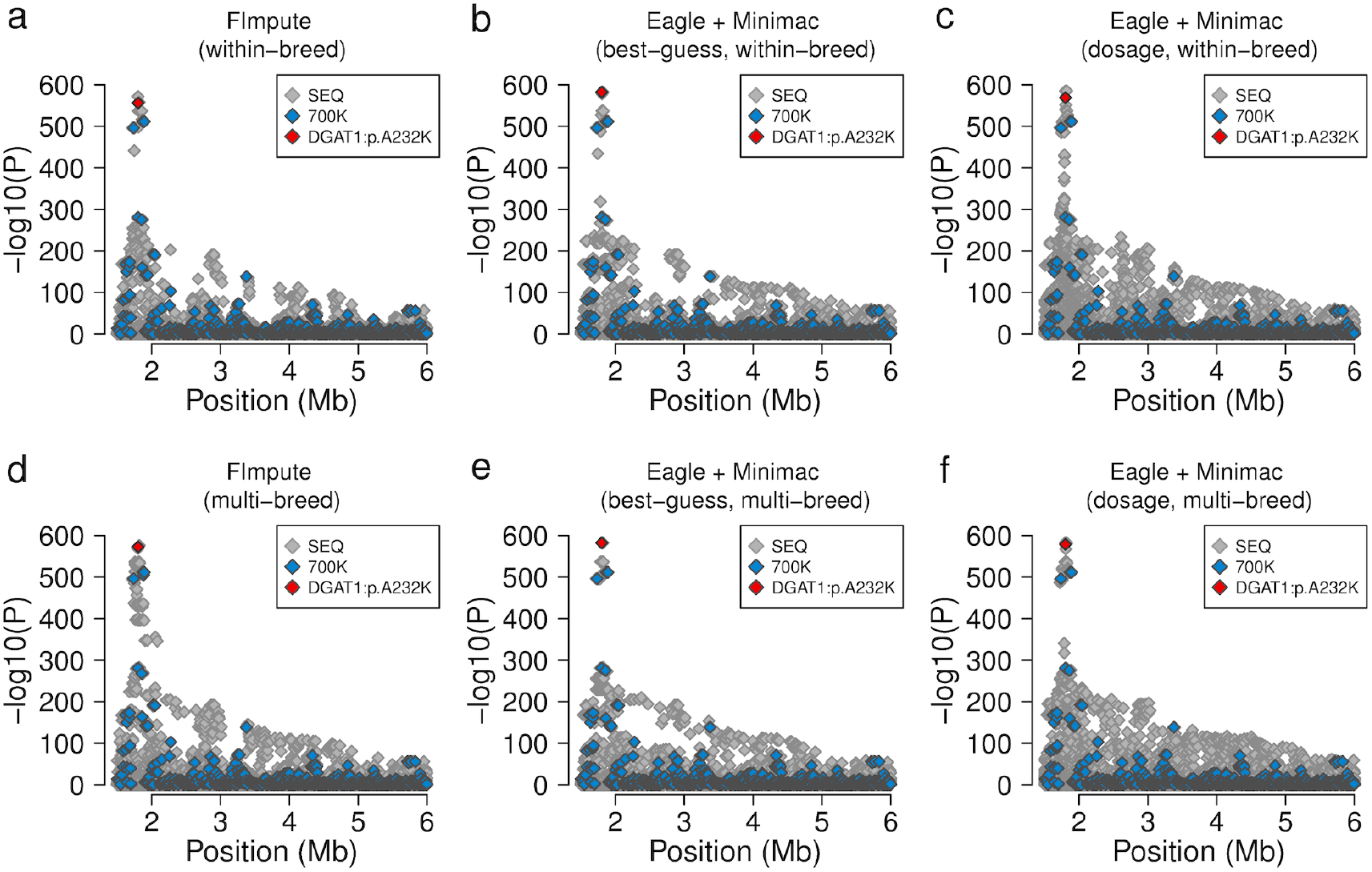
Fine-mapping of a fat percentage QTL on bovine chromosome 14. (a-e) Association between 48,641 imputed sequence variants located at the proximal end of bovine chromosome 14 and milk fat percentage in 6958 Fleckvieh animals. Genotypes for the association studies were imputed using either *FImpute* (a, d) or *Minimac* (b, c, e, f) with either within-(a-c) or multi-breed reference populations (d-f). Grey and blue colours represent sequence and array-derived variants. The red symbol represents the p.A232K-mutation in the *DGAT1* gene.

The imputation algorithm and composition of the reference population had a large effect on the ability to detect an association between the GHR:p.F279Y-variant and milk fat percentage (Figure 3). The GHR:p.F279Y-variant was not significantly associated (P>1×10^−8^) when the genotypes were imputed using *FImpute* or *Minimac*_BG_ and a within-breed reference population reflecting high imputation error rates (Table 3). There were 350 and 395 variants detected, respectively, that had lower P values than rs385640152. Association tests with genotypes that were obtained using *Minimac*_DOS_ (within-breed) or *FImpute* (multi-breed) revealed significant association of rs385640152 (P=1.0×10^−15^, 3.0×10^−10^). However, six and 1089 variants had lower P values than rs385640152. When the genotypes were inferred using *Minimac* and a multi-breed reference population, rs385640152 was the most significantly associated variant reflecting higher accuracy of imputation (Figure 3e,f, Table 3).

**Figure 3.**
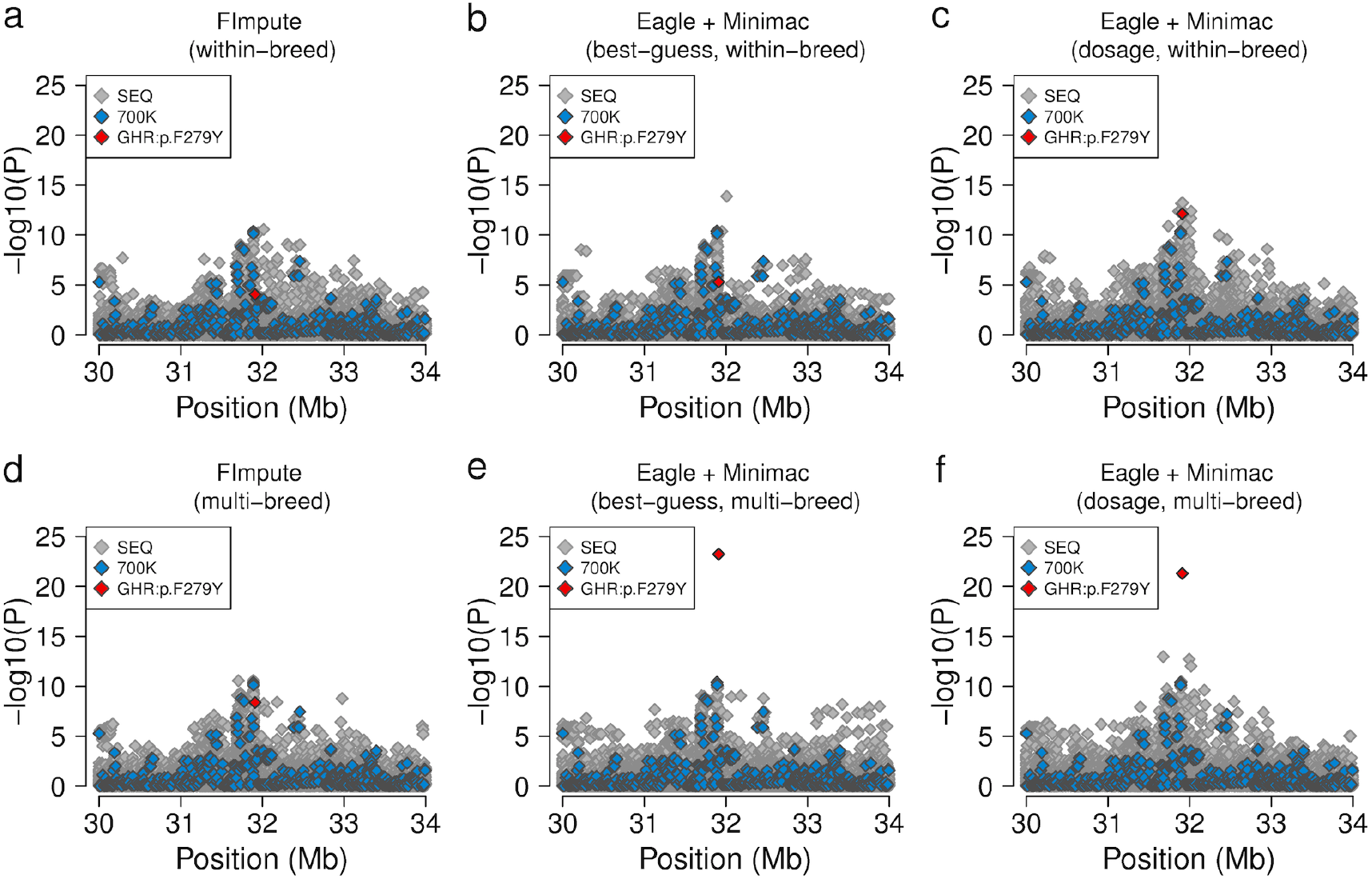
Fine-mapping of a fat percentage QTL on bovine chromosome 20. (a-e) Association between 29,205 imputed sequence variants on bovine chromosome 20 and milk fat percentage in 6958 Fleckvieh animals. Genotypes for the association studies were imputed using either *FImpute* (a, d) or *Minimac* (b, c, e, f) with either (ac) or multi breed reference populations (d-f). Grey and blue colours represent sequence and array-derived variants, respectively. The red symbol represents the p.F279Y-mutation in the *GHR* gene.

### Fine-mapping of five fat percentage QTL with imputed sequence variants

The fine-mapping of two known QTL indicated that genotypes that are imputed using *Minimac* and considering a multi-breed reference population are an accurate source for association tests, particularly when predicted allele dosages rather than best-guess genotypes are used as explanatory variables. We imputed 23,256,743 sequence variants in 6958 FV animals using *Minimac* and a multi-breed reference population and performed association tests between imputed allele dosages and fat percentage. The mean and median accuracy of imputation for 23,256,743 sequence variants (r^2^-values from *Minimac*) was 0.71 and 0.95, respectively (see **Additional File 4 Figure S4**). 78.3 and 73.2% of the imputed sequence variants had r^2^-values greater than 0.3 and 0.5, respectively. Our association study revealed seven genomic regions with significantly associated variants (*i.e*., QTL), including two QTL on BTA14 and BTA20 that encompassed the *DGAT1* and *GHR* genes (Figure 4a, Figure 2f, Figure 3f). The top variants were imputed sequence variants at all QTL.

**Figure 4.**
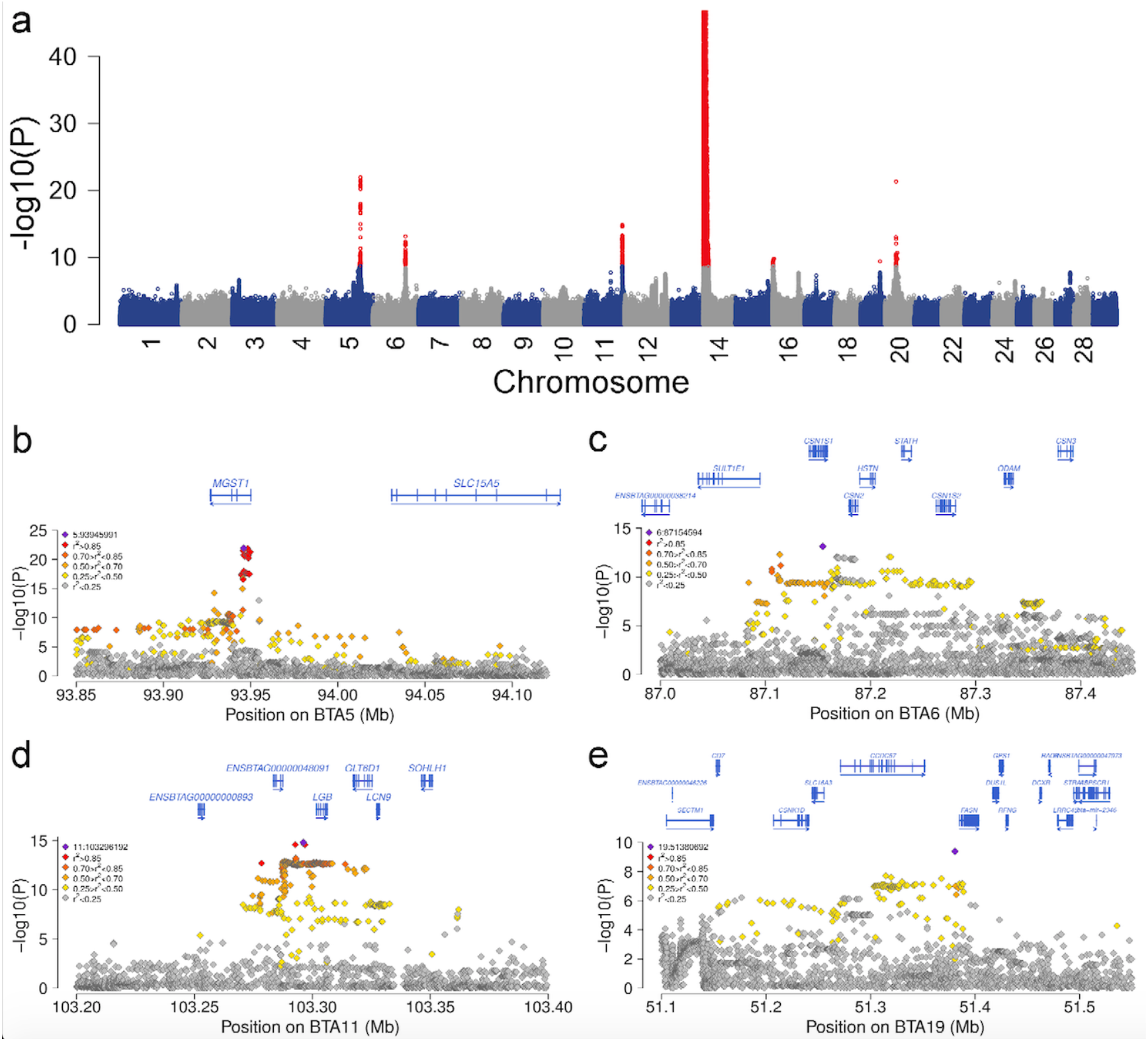
Fine-mapping of fat percentage QTL in Fleckvieh cattle. (a) Manhattan plot representing the association of 23,256,743 imputed sequence variants with milk fat percentage in 6958 Fleckvieh animals. Red colours represent significantly associated variants (P<2.15×10^−9^). The y-axis is truncated at −log10(10^−4^°). (b-e) Fine-mapping of four QTL for milk fat percentage on bovine chromosomes 5, 6, 11 and 19. Different colours represent the linkage disequilibrium between the most significantly associated variant (violet) and all other variants. Blue arrows indicate the direction of the gene transcription.

A total of 239 variants located between 91,857,670 and 93,955,207 bp on BTA5 were significantly associated (P<2.1×10^−9^) with fat percentage (Figure 4b, see **Additional file 9 Table S1**). Thirteen variants in the first intron of the *MGST1* gene *(microsomal glutathione S-transferase 1)* were in high LD (r^2^>0.95) with each other and had markedly lower P values (P<6.5×10^−21^) than all other variants. These 13 variants were also associated with milk composition in New Zealand dairy cows [23]. The top variant (rs208248675) was located at 93,945,991 bp. The rs208248675 A-allele had a frequency of 19.4% in FV cattle and it decreased fat percentage.

A QTL for fat percentage on BTA6 encompassed the casein gene complex (Figure 4c, see **Additional file 9 Table S1**). One hundred and sixty-four variants with P<2.1×10^−9^ were located between 87,084,144 and 87,296,017 bp. The most significantly associated variant (rs109193501 at 87,154,594 bp, P=7.6×10^−14^) was located in an intron of the *CSN1S1* gene *(casein alpha s1)*.

Two hundred and sixty-two variants located within a 50 kb interval (between 103,274,736 and 103,324,728 bp) on BTA11 were associated with fat percentage (Figure 4d, see **Additional file 9 Table S1**). The top variant (rs381989107 at 103,296,192 bp, P=1.5×10^−15^) was located 5472 bp upstream of the translation start site of the *LGB* gene *(beta-lactoglobulin*, also known as *progestagen associated endometrial protein (PAEP))*. Two missense variants (p.G80D and p.V134A) in the *LGB* gene (rs110066229 at 103,303,475 bp and rs109625649 at 103,304,757 bp) that distinguish the LGB protein variants A and B [42] were in high LD with rs381989107 and had P values less than 4.5×10^−12^ (see **Additional file 9 Table S1**). A variant in the promoter of the *LGB* gene that causes an aberrant LGB expression in Brown Swiss cattle (OMIA 001437-9913) [43] was not polymorphic in the sequenced FV animals.

A QTL on BTA16 included 14 significantly associated variants located between 481,671 and 3,265,708 bp (see **Additional file 9 Table S1**). The top association signal (rs380194132, P=1.6×10^−10^) resulted from an intergenic variant that was located at 3,242,688 bp. Since the genomic region that included associated sequence variants with similar P values extended over almost 3 Mb, the identification of putative candidate gene(s) underlying the QTL was not attempted (see **Additional file 5 Figure S5**).

On BTA19, a single variant located at 51,380,692 bp was significantly associated (rs208289132, P=4.1×10^−10^) with fat percentage (Figure 4e, see **Additional File 9 Table S1**). The associated variant was located 4230 bp upstream of the translation start site of the *FASN* gene *(fatty acid synthase*).

### Computing resources required

We assessed the computing resources required to impute genotypes at 23,256,743 sequence variants for 6958 target animals using 1577 sequenced animals as a reference. *Eagle* and *Minimac* were run on 10 processors per chromosome, whereas *FImpute* was run on a single processor. *FImpute* ran out of memory and failed to impute genotypes for chromosomes 12 and 23 likely because of large structural variants located on both chromosomes (see **Additional file 6 Figure S6**). The process (CPU) time to infer haplotypes and impute sequence variants for 27 chromosomes was 1037, 490 and 146 hours, respectively, for *Eagle, Minimac* and *FImpute*.

The wall-clock time and RAM usage for *FImpute* was between 2.9 and 9.6 hours and 6.7 and 29.7 gigabytes per chromosome, respectively (see **Additional file 7 Figure S7**). The estimation of haplotype phases took between 1.6 and 5.9 hours using *Eagle* and it required between 1.8 and 3.4 gigabytes of RAM with between 11,178 and 38,009 SNPs per chromosome for 6958 animals. The imputation of sequence variants using *Minimac* took between 2.5 and 9.4 hours and it required between 6.3 and 21 gigabytes of RAM per chromosome.

The estimation of haplotype phases for 1577 reference animals using *Eagle* (multithreaded on 10 CPUs) took between 4.2 and 15.1 hours with between 413,371 and 1,444,299 sequence variants, respectively, and it required between 15.4 and 51.7 gigabytes of RAM per chromosome.

## Discussion

We evaluated the accuracy of imputation from dense genotypes to sequence variants using a population of 1577 sequenced animals which is a four-to eight-fold increase in sample size compared to previous studies in cattle [2, 17, 18]. Our findings are likely to be relevant for many cattle populations because we considered animals from two breeds with different effective population sizes as validation panel and inferred sequence variants on six chromosomes that reflect a broad spectrum of LD [44]. The accuracy of imputation was the correlation between imputed and sequenced genotypes (r_IMP,SEQ_), which is a measure that depends less on the allele frequencies of the sequence variants than *e.g*., the proportion of correctly imputed genotypes [45]. However, the cross-validation procedure that was applied in our study yields reliable results only when the quality of the sequence data is high. Low fold sequencing coverage and sequencing or assembly errors may result in flawed genotypes [46]. When genotypes at such positions are masked and subsequently imputed, genotype imputation might provide true genotypes that differ from the sequenced genotypes thereby underestimating the accuracy of imputation. Considering that the genotype error rates were low in the sequenced FV and HOL animals [2, 47], it is unlikely that our findings are biased due to flawed genotypes at some sequence variants.

The r_IMP,SEQ_-values varied considerably across chromosomes and dropped at several positions along the genome. Previous studies using array-derived genotypes showed that a decline in imputation accuracy may indicate intra-or inter-chromosomal misplacement of SNPs [12, 48, 49]. We detected low r_IMP,SEQ_-values at regions where the bovine genome contains large segmental duplications [40, 41]. The population-wide imputation of sequence variants for chromosomes 12 and 23 was not possible using *Fimpute*. Assessing the accuracy of imputation along these chromosomes using cross-validation revealed large segments with low imputation quality on both chromosomes including the highly variable major histocompatibility complex [50]. We were able to eventually impute genotypes for BTA12 and BTA23 using *FImpute* when those segments were excluded from the reference panel. Imputing genotypes within both segments was possible using *Minimac*. However, the inferred genotypes are flawed and must be treated with caution as our results show (see **Additional file 6 Figure S6**). While an improved assembly of the reference genome may resolve some of these problems, a better resolution of large structural elements is not possible using short paired-end DNA sequencing [51]. Since flawed genotypes compromise downstream analyses [24, 25], it is advisable to exclude sequence variants within segments of low imputation quality from the target population and retain the array-derived genotypes only.

We assessed the performance of *FImpute* and *Minimac* because both methods can impute millions of polymorphic sites in large populations within a reasonable time. Previous studies showed that *FImpute* is more accurate than other tools with similar computing costs [52, 53]. The population-based genotype imputation algorithm implemented in *Minimac* is highly accurate and fast because it takes previously phased genotypes as input [11, 54]. We phased the target population using *Eagle* because it enables timely and accurate haplotype inference [32]. Although the computing costs to impute 23 million sequence variants in 6958 animals were more than ten-fold higher with *Eagle* and *Minimac* than *FImpute*, the wall-clock time was only 1.67-fold higher because *Eagle* and *Minimac* enable multi-threading to reduce the process time [32, 54]. The difference in computing costs is greater when haplotypes are not available for the reference animals. Since many variant detection pipelines include phasing and imputation algorithms [2, 47, 55], the computing time required to phase the sequence data was not considered in our study. However, the phasing of the reference panel was necessary because the variant detection pipeline of the 1000 bull genomes project was applied to successive 5 Mb segments rather than whole chromosomes [2].

The accuracy of imputation was higher with *Minimac* than *FImpute*, although *Minimac* does not consider pedigree information. Most individuals of the 1000 bull genomes population were selected in a way that they represent a large proportion of the genetic variation of current populations [2, 56]. They are likely less related with each other than a random subset from the population, although some parent-offspring pairs were included. The accuracy of imputation might be higher when reference and target animals are closely related particularly when *FImpute* is used [53]. However, dense genotypes for reference and target animals, *e.g*., sequence and HD-derived genotypes, can be used to identify short shared haplotypes among apparently unrelated animals that originate from ancient ancestors possibly predating the separation of breeds [9, 57]. Including reference animals from various breeds increased the accuracy of imputation, particularly in FV cattle. The principal components analysis revealed that animals from many breeds clustered nearby the FV population indicating that they are distantly related whereas the HOL animals formed a cluster that was separated from all other breeds. The benefit of a multi-breed reference population on the accuracy of imputation was less pronounced in HOL cattle. Since the number of reference animals was nearly twice as high in HOL, our results may also corroborate the suggestion that multi-breed reference panels are particularly useful to impute genotypes when the reference population is small [18, 58, 59]. In agreement with previous studies in cattle [2, 18], a multi-breed reference population increased the accuracy of imputation at rare variants likely reflecting the shared ancestry of many cattle populations and the limited number of sequenced animals per breed. However, it seems advisable to periodically evaluate different imputation strategies because multi-breed reference populations can also compromise the imputation of sequence variants when the diversity of the reference panel increases [26].

The population-based imputation of sequence variants was particularly advantageous at variants with low MAF corroborating previous findings [12, 53, 60]. The true genotypes were more correlated to predicted allele dosages than best-guess genotypes which is in line with the findings of Brøndum et al. [18]. Moreover, our results show that more sequence variants are polymorphic in the target population when allele dosages are considered instead of best-guess genotypes. An association study between allele dosages and milk fat percentage revealed that two variants with a MAF less than 10% in the *DGAT1* [34, 35] and *GHR* [36] genes were the most significantly associated variants at two QTL on BTA14 and BTA20. This finding demonstrates that utilizing predicted allele dosages in association studies with imputed sequence variants can pinpoint causal mutations which agrees with Zheng at al. [61] and Khatkar et al. [62]. So far, imputed sequence variants did not substantially improve the accuracies of genomic predictions in real data which might partly result from flawed genotypes at low-frequency variants [17, 63]. We show that allele dosages are more accurate than best-guess genotypes particularly at variants with low MAF because they better reflect imputation uncertainty. However, the analysis of allele dosages for millions of sequence variants in tens of thousands of individuals for genomic predictions is computationally costly and has not been attempted so far [17, 64]. Since the proportion of variants with low MAF is more than three-fold higher in sequence than HD variants, further research is warranted to investigate if allele dosages may enhance genomic predictions with imputed sequence variants.

Our findings show that the composition of the reference population and the choice of the imputation algorithm are critical to infer accurate genotypes thereby enabling us to pinpoint causal mutations in association studies with imputed sequence variants. When the accuracy of imputation is low, causal variants may be “buried” among other variants in LD (*e.g*., Figure 3a,d). In such situations, association testing will not reveal true causal mutations because variants with lower P values are likely to be prioritized as candidate causal variants [23]. However, true causal variants are not necessarily the top variants in association studies even when their genotypes are almost perfectly imputed. Although the imputation accuracy for the causal p.A232K-polymorphism in the *DGAT1* gene was greater than 99.5%, several adjacent variants had identical or slightly lower P values likely indicating sampling errors [65] or synthetic associations [66, 67]. Nevertheless, the mutations in the *DGAT1* and *GHR* genes offer a convenient approach to evaluate the ability to detect causal mutations with imputed sequence variants in cattle, because both variants segregate in many breeds and phenotypes for fat percentage are readily available.

Our association study with more than 23 million sequence variants detected seven QTL that were significantly associated with milk fat percentage. Applying a less stringent significance threshold [68] would reveal additional QTL (see **Additional file 8 Figure S8**). Since we were able to pinpoint the causal mutations at two QTL, it is likely that causal mutations for other QTL are among the most significantly associated variants. However, the QTL identified in our study include several sequence variants in high LD and with similar P values rendering the identification of causal sites difficult. Association studies in animals from multiple breeds may facilitate differentiation between causal and non-causal sites in LD [2]. Causal sites may be located on ancient haplotypes that still segregate across multiple breeds as is the case for the fat percentage QTL on BTA5. Our association study revealed thirteen candidate causal variants in high LD in an intron of the *MGST1* gene that had markedly lower P values than all other variants. It is very likely that one of those variants is the actual causal polymorphism because they were also highly significantly associated with milk composition in another dairy cattle breed [23]. An improved functional annotation of the bovine genome and a fine-mapping strategy that prioritizes sequence variants using functional information may facilitate to pinpoint causal mutations at such QTL [22, 69].

## Conclusions

Genotypes for millions of sequence variants can be accurately inferred in large bovine cohorts using population-based imputation algorithms. The accuracy of imputation for variants with low MAF is higher with multi-breed reference populations. The accuracy of imputation is not constant across chromosomes and may drop at regions where the genome contains large segmental duplications. Considering predicted allele dosages rather than best-guess genotypes as explanatory variables is beneficial for association studies with imputed sequence variants. The composition of the reference population and the choice of the approach used to infer genotypes are critical to detect causal mutations in association studies with imputed sequence variants.

## Declarations

### Ethics approval and consent to participate

Not applicable

### Consent to publication

Not applicable

### Availability of data and material

All relevant data are included in the manuscript and its supporting files

## Competing interests

The authors declare that they have no competing interests.

## Funding

HP was financially supported by a Deutsche Forschungsgemeinschaft (DFG) postdoctoral fellowship.

## Authors’ contributions

HP, IMM, HDD and MEG designed the experiments, HP analyzed the data, PJB provided support in computing, RF and RE provided genotype and phenotype data, HP wrote the manuscript. All authors read and approved the final manuscript.

## Acknowledgements

We thank Arbeitsgemeinschaft Süddeutscher Rinderzüchter und Besamungsorganisationen e.V. (ASR), Arbeitsgemeinschaft österreichischer Fleckviehzüchter (AGÖF) and Fürderverein Biookonomieforschung e.V. (FBF) for providing genotyping data. We acknowledge the 1000 bull genomes project for providing sequence variants.

## Additional files

**Additional file 1 Figure S1**

File format: tif

Title: **Principal component analysis in 1577 sequenced animals.**

Description: (a-c) Plot of the top two principal components of the genomic relationship matrix. Different colours and symbols represent different breeds. The partners of the 1000 bull genomes consortium assigned the animals to breeds (b) Non-grey symbols indicate 249 animals that were considered as Fleckvieh animals. (c) Green symbols indicate 450 Holstein animals.

**Additional file 2 Figure S2**

File format: tif

Title: **Sequence variants that are invariant in the validation populations.**

Description: (a-d) Proportion of sequence variants that were invariant after imputation in the FV (a, c) and HOL (b, d) validation populations for six chromosomes analysed and different MAF classes. (e, f) The correlation between imputed and true genotypes was calculated using only sequence variants that were polymorphic in the FV (e) and HOL (f) validation populations.

**Additional file 3 Figure S3**

File format: tif

Title: **Imputation accuracy along six chromosomes in Fleckvieh cattle.**

Description: (a-f) The correlation between true and imputed genotypes for sequence variants located within successive 1 Mb windows on six chromosomes. Different colours and symbols represent correlation coefficients obtained using different imputation scenarios. *Insets*: HD SNP coverage and sequence variant density along the chromosome. Black and red colours represent the number of SNPs that were included in the BovineHD Bead Chip (HD) and sequence (Seq) variants (x1000, (K)) that were polymorphic in the multi-breed reference population, respectively, per million basepairs (Mb).

**Additional file 4 Figure S4**

File format: tif

Title: **Accuracy of imputation for 23,256,743 sequence variants in 6958 animals.**

Description: The mean (red) and median (blue) accuracy of imputation (r^2^-values were obtained using *Minimac*) for 23,256,743 sequence variants (grey bars) as a function of the minor allele frequency. The dotted lines represent the mean and median accuracy of imputation across all sequence variants.

**Additional file 5 Figure S5**

File format: tif

Title: **Detailed view of a milk fat percentage QTL on bovine chromosome 16**

Description: Different colour represents the linkage disequilibrium between the most significantly associated variant (violet) and all other variants. Blue arrows indicate the direction of the gene transcription.

**Additional file 6 Figure S6**

File format: tif

Title: **Imputation accuracy along chromosomes 12 and 23 in Fleckvieh cattle.**

Description: (a-b) The correlation between true and imputed genotypes for sequence variants located within successive 1 Mb windows on chromosomes 12 and 23. Different colours and symbols represent correlation coefficients obtained using different imputation scenarios. (c-d) HD SNP coverage and sequence variant density along chromosomes 12 and 23. Black and red colours represent the number of SNPs that were included in the BovineHD Bead Chip (HD) and sequence (Seq) variants (x1000 (K)) that were polymorphic in the multi-breed reference population, respectively, per million basepairs (Mb). We were eventually able to impute sequence variants for BTA12 and BTA23 using *FImpute* when we discarded sequence variants that were located between 70 and 77 Mb and between 25 and 30 Mb, respectively, from the reference panel.

**Additional file 7 Figure S7**

File format: tif

Title: **Computing resources required to impute 23,256,742 sequence variants in 6958 animals.**

Description: The wall-clock times (a) and random-access memory (RAM) usage (b) required to infer haplotypes and genotypes with *FImpute* (green), Eagle (dark blue) and *Minimac* (light blue) were assessed on 12-core Intel^®^ Xeon^®^ processors rated at 2.93 Ghz with 96 gigabytes of RAM. *FImpute* ran out of memory and did not finish when we attempted to infer genotypes for BTA12 and BTA23. *FImpute* was ran on a single processor whereas *Eagle* and *Minimac* used 10 processors per chromosome.

**Additional file 8 Figure S8**

File format: tif

Title: **Detailed view of a milk fat percentage QTL on bovine chromosome 27**

Description: Different colour represents the linkage disequilibrium between the top variant (violet) and all other variants. Blue arrows indicate the direction of the gene transcription. The top variant (36,211,258 bp) was associated with fat percentage (P=1.9×10^−8^) albeit not at genome-scale. Twenty-two variants in high LD (r^2^>0.68) with the top variant were located between 36,200,888 and 36,253,406 bp and had P values less than 7.7×10^−7^. Among those were three candidate causal variants (36,211,252 bp with P=2.4×10^−8^, 36,211,708 bp with P=2.8×10^−8^, 36,209,319 bp with P=3.3×10^−8^) for fat content in the early lactation that were reported in Daetwyler *et al*. [2].

**Additional file 9 Table S1**

File format: tif

Title: **Significantly associated variants at five fat percentage QTL**

Description: Variants located on chromosomes 5, 6, 11, 16 and 19 with P values less than 2.1×10^−9^. The positions of the variants correspond to the UMD3.1 assembly of the bovine genome. The substitution effects (beta, standard error of beta) are given for the alternative allele. The R2 value is the estimated accuracy of imputation from *Minimac*.

